# Variation in the response to malaria in Diversity Outbred mice

**DOI:** 10.1101/2021.09.29.462483

**Authors:** Avni S. Gupta, Victoria Chevée, Adam S. Kirosingh, Nicole M. Davis, David S. Schneider

## Abstract

We infected Diversity Outbred mice with *Plasmodium chabaudi* to better understand how the host response to infection can vary and to try to identify genetic loci responsible for this variation. We identified two loci correlating with binary traits: one on chromosome two was linked to undetectable parasite loads and another on chromosome ten which was linked to death. Though we tested many variable traits, none of those reached statistical significance using the 489 mice we tested.

## Introduction

Hosts can vary in their response to a single pathogen. For example, *P. falciparum*, is one of five parasites that cause human malaria and this species alone can cause different outcomes that include anemia, placental infections, lung, kidney, liver and brain dysfunction. Some of this is due to parasite variation, but much of it is due to differences in host responses. We would like to understand the available landscape of variation and then determine which aspects of this landscape are under host genetic control.

Malaria is a life-threatening infectious disease which, although preventable and treatable, still affected an estimated 228 million people worldwide in 2018, leading to 405,000 deaths (World Health Organization). Host genetics are an important modifier of malaria severity (Tran and Crompton 2020) as humans have evolved protective resistance and disease tolerance mechanisms to maintain health in the face of infections. Examples of evolved host defenses to malaria include mutations in hemoglobin subunit beta (sickle cell trait), alpha-thalassemia, glucose-6-phosphate dehydrogenase (G6PD), and nitric oxide synthase 2 (Driss *et al.* 2011; Kariuki and Williams 2020).

To understand the available landscape of variation, we need to follow disease symptoms and signs over the course of an infection so that we can measure maximal and minimal levels of these phenotypes as well as rates of change. This is very difficult to do with human patients. As a substitute, we chose to study a mouse model of malaria because malaria is an important human disease and the mouse model lets us ask some basic questions about disease ecology in a laboratory setting, where we can monitor the mice daily. Like humans, inbred laboratory mouse strains vary in susceptibility to *Plasmodium* (Stevenson *et al.* 1982; Foote *et al.* 1997; Bopp *et al.* 2010; Laroque *et al.* 2012). The murine strain *Plasmodium chabaudi chabaudi* causes an array of symptoms similar to those produced in human malaria *(Stephens et al.* 2012), differentially affecting temperature, anemia, liver and kidney function. Variation in susceptibility to *P. chabaudi* has been used to identify ten *P. chabaudi* resistance (*Char*) loci in mice. These *Char* loci were mapped by characterizing the response to infection in mice derived from crosses between a susceptible strain (A/J, C3H/He, or SJL/J) and a resistant strain (C57BL/6J). Parasitemia (percent of infected red blood cells (RBC)), survival, erythropoiesis, and parasite growth rate (Huang *et al.* 2018) were used to map single nucleotide polymorphisms (SNPs)s that correlated with disease outcomes.

One strain of mice doesn’t tell you how mice behave, and no two strains represent the entire diversity of the species. New genetic tools now allow researchers to leverage the genetic diversity of the mouse species as a whole to identify drivers of disease. The Diversity Outbred (DO) mouse population, created and maintained by The Jackson Laboratory (JAX), is the product of an eight-parent breeding design that covers nearly 90% of the most common mouse genetic variants (Roberts *et al.* 2007). DO mice are derived from five inbred strains (C57BL/6, 129S1/SvImJ, NOD/ShiLtJ, NZO/HILtJ, A/J) and three wild-derived strains (PWK/PhJ, WSB/EiJ, CAST/EiJ). (Churchill *et al.* 2004; Roberts *et al.* 2007). The DO population can be used in at least two ways to study the genetic contribution to a phenotype. First, the population will exhibit a broader range of phenotypes rather than the relatively uniform responses seen in one inbred mouse line. This can help identify phenotypes that better represent the diversity of human disease that might otherwise be missed by focusing on your favorite mouse strain. Populations like this help remind us that “the mouse” doesn’t behave in one way, but has diverse responses in the same way that humans do. Second, these diverse mice can be used for genetic mapping. An advantage of this collection over a true wild population is the high frequency of minor alleles (12.5%) from each parental strain, which produces a relatively high number of homozygotes in a small population of mice. (*Churchill et al.* 2012).

Here, we exploited host genetic variation in the DO mice to measure the range of phenotypes produced by *Plasmodium-*infected mice, and to map loci controlling this range of responses. We infected 489 DO mice with *P. chabaudi* AJ and measured changes in parasite loads, temperature, weight, RBC counts from early in the infection until peak pathology. This let us see which health measures varied and might serve as phenotypes in a GWAS. We used these data to cluster the mouse phenotypes and determined phenotype linkage to genetic loci using QTL analysis with a high resolution genotyping array containing 616,136 SNPs. In the end, two loci showing significant correlations with binary phenotypes: The first is a locus that mapped a negative blood smear phenotype to a 0.6Mb interval on chromosome 2 containing two genes (*Itga4,* an integrin subunit, and *Cerkl,* a ceramide kinase-like protein). The second, locus a 2.38Mb interval on chromosome 10, was associated with death. This locus contains 20 genes with known SNP variants among the eight parental strains, including *Foxo3,* an immune modulator in monocytes.

## Materials and Methods

### Mice

Female mice were purchased from The Jackson Laboratory (JAX) (Bar Harbor, Maine, USA) and maintained in the Stanford University animal facility. Mouse strains used in this study are listed in Table S1. All experiments were performed in accordance with institutional guidelines and approved protocol (APLAC-30923). The first batch of DO mice was from Generation 23, Litter 2. The last batch of DO mice was from Generation 30, Litter 1. Mice of each parental strain were included in experiments for genotyping controls and infection phenotype analysis (N=4-7 for each parental strain).

All mice were acclimated in cohoused cages of 5 animals in the Stanford University animal facility for 7–10 days prior to being used in experiments. Female mice of a similar age (8-15 weeks old) were infected with *Plasmodium chabaudi chabaudi* AJ (Malaria Research and Reference Reagent Resource Center [MR4]) to limit variance in the murine response to this parasite caused by sexual dimorphism and age (Stephens *et al.* 2012). Upon recommendation by JAX, the DO mice were not shuffled to randomize microbiota on arrival to the facility due to concerns of aggression towards unfamiliar cagemates.

### Temperature Probes

Mice were anesthetized locally with 2% lidocaine solution (100 μg delivered per dose) and implanted subcutaneously with electronic temperature and ID transponders (IPTT-300 transponders, Bio Medic Data System, Inc) one week prior to *P. chabaudi* infection. Temperatures were recorded using a DAS-7006/7s reader (Bio Medic Data System, Inc).

### Infection

C57BL/6 female mice were given intraperitoneal injections of 100 μL of thawed *P. chabaudi chabaudi* AJ-infected RBCs (iRBCs) from the parasite stock described previously. In order to monitor parasitemia, thin blood smears were prepared from tail blood, methanol fixed, and Giemsa stained (Gibco-KaryoMAX). Parasitemia was quantified under light microscopy at 100x magnification daily until parasitemia reached 10-20% (9 days post-infection). Then experimental mice were injected intraperitoneally with 100 μL containing 10^5 freshly obtained iRBCs diluted in Krebs saline with glucose (KSG).

### Longitudinal infection monitoring

Weight and temperature were recorded daily between 9 AM and 12 PM during the experiment. In situations where mice were not implanted with temperature probes, the temperature was determined using a rectal thermometer (BAT-12, Physitemp Instruments Inc.). Approximately 19 μL of blood was collected daily via tail nicking of restrained mice using sterilized surgical scissors. Sodium acetate warmers were used on occasion to increase tail blood flow. Blood was collected into EDTA-coated 50 μL capillary tubes to inhibit clotting. Beginning on day five, roughly 2 μL of blood per animal was used to generate thin blood smears, which were subsequently prepared and counted as detailed above to measure parasitemia. Each day, 2 μL of blood was diluted in one mL of Hanks’ Balanced Salt Solution (HBSS) to count the number of RBCs. Absolute RBC counts were obtained on an Accuri C6 Flow Cytometer (BD Biosciences) using forward and side scatter.

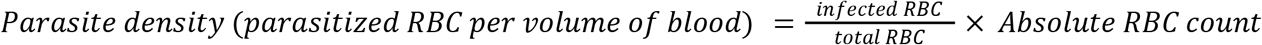

### Genotyping

Tail clippings (2 mm) were collected on Day 0 of the infection, flash-frozen in liquid nitrogen and stored at −80°C for later processing. DNA was extracted from the tail clippings via overnight incubation in a phenol-chloroform based extraction buffer (50 mM Tris-HCl pH 8.0, 100 mM EDTA pH 8.0, 100 mM NaCl, 1% SDS) followed by two ethanol washes. Samples were quantified on Nanodrop to ensure quality and quantity. DNA samples were genotyped at Life Technologies (Santa Clara, CA) on a custom Axiom Mouse Non-Human Genotyping Array with 616,136 SNPs distributed throughout the mouse genome with an average spacing of 4.3 kb, allowing for a high resolution analysis. This array is annotated using the mm10 mouse build (GRCm38-Genome Reference Consortium Mouse Build 38). Samples were analyzed in plates of 96. Replicates of each of the founder strains were added to the genotyping array in order to determine founder strain genotypes at each of the SNPs for future haplotype reconstruction.

### Genetic linkage mapping and clustering

Primary quality control analysis of the compiled genotyping data from all samples was completed by the Life Technologies Bioinformatics Services Team on Axiom Analysis Suite. Axiom Analysis Suite was then used to export raw call codes for SNPs that passed the quality thresholds. Qtl2 (R package by Karl Broman) was used to reconstruct haplotype states for each of the individual DO animals using a Hidden Markov Model (HMM). The HMM estimates the haplotype contribution of each of the eight founder strains at each SNP to each mouse by summing the probabilities contributed by each founder. A mixed model was fit with sex and experimental group as additive covariates and a random effect was included to account for kinship. We then performed QTL analysis to determine loci of interest on the basis of logarithm of the odds (LOD) scores (Broman *et al.* 2019). LOD scores measure the likelihood of genetic linkage: the higher the LOD score, the more likely that the locus identified is linked to the phenotype queried. Significance thresholds were determined by analyzing 1,000 permutations of the genome scan with randomization of the phenotype data among individuals. We determined 1.5-LOD support intervals for significant peaks. This package was linked to an SQLite database using the mm10 mouse build to determine genes in the identified intervals (https://figshare.com/articles/SQLite_database_with_mouse_gene_annotations_from_Mouse_Genome_Informatics_MGI_at_The_Jackson_Laboratory/5280238/6).

## Results

We had two goals in measuring the response of diverse mice to *Plasmodium*: First, we wanted to determine at a species level, the extent of variation of responses to an infection under laboratory conditions and to identify correlations between different phenotypes. Second, we wanted to determine which of these variations could be mapped using GWAS so that we could identify the underlying genetic factors that determine this variation.

To capture most of acute infection and disease, we conducted a longitudinal study challenging DO mice with *P. chabaudi* and monitoring them regularly post infection. We used a dose of *P. chabaudi* AJ that kills approximately 20% of infected C57BL/6 mice, and thus we anticipated finding both more and less sensitive strains of mice (Torres et al. 2016). Each day of the infection we measured the following signs of disease: weight, temperature, RBC concentration, parasitemia (proportion of infected RBC per total RBC), and parasite density (10^6^ parasites per microliter of blood). We examined mice on day 0, as an internal uninfected control for each mouse, and on days 5 through 15, as peak pathology occurs during this interval.

To determine the range of phenotypes resulting from *P. chabaudi* infections, we tested the 8 founder strains of the DOsto determine the limits and timing of phenotypes. We found a broad range of results: Some mouse strains showed high lethality when infected under this regime while others suffered essentially no disease(Figure 1A-E). For example, the WSB/EiJ mice did not lose weight, become hypothermic, or become severely anemic when infected with *P.chabaudi* AJ and their infection occurred later than the other mice. C57BL/6 mice had mid-range phenotypes with about half of the other mouse strains doing better or worse. The DO mice fit into this continuous range of phenotypes (Figure 1F-J).

**Figure 1.**
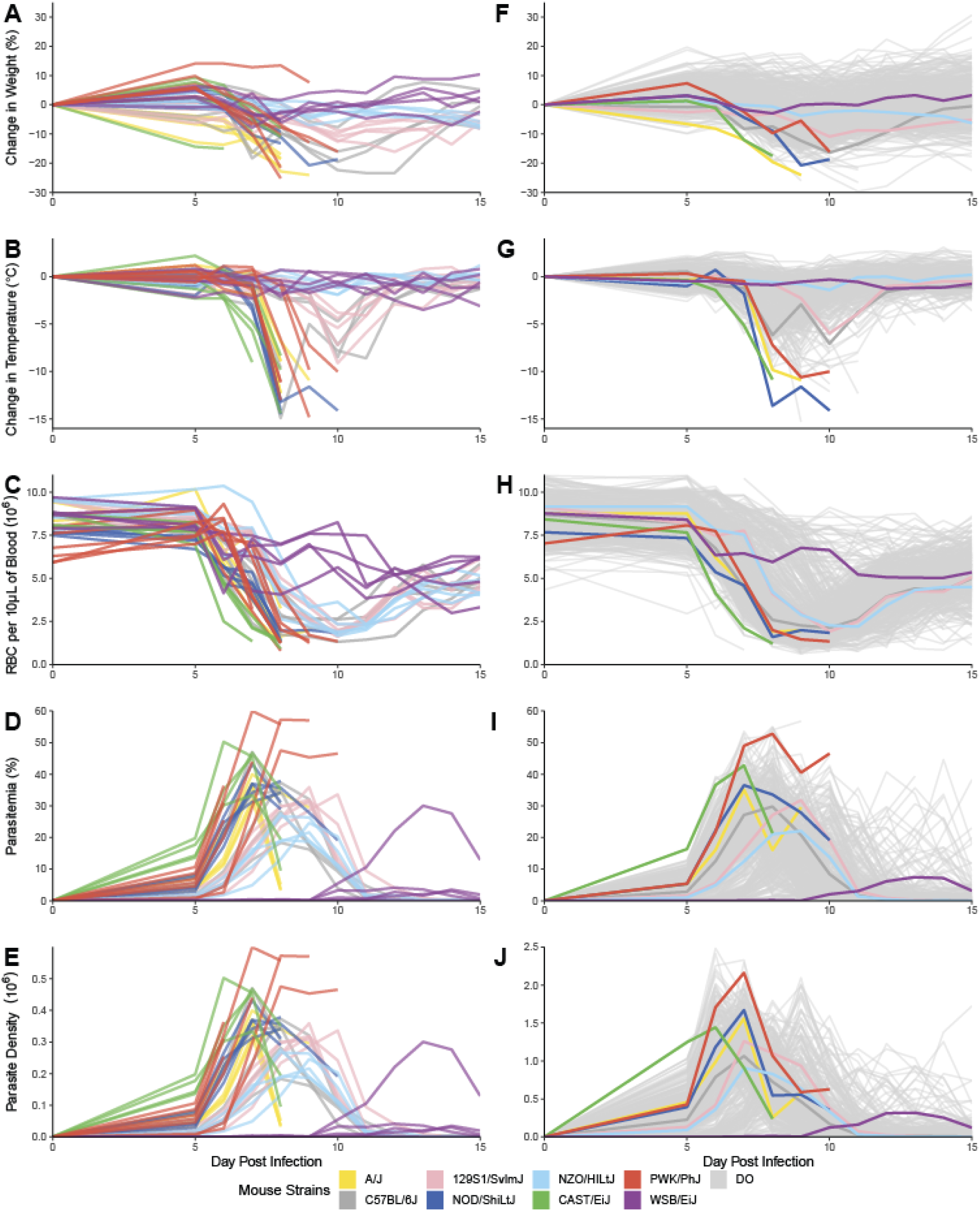
Longitudinal infection monitoring for parental mouse strains and DO mice reveals phenotypic variability There is phenotypic variability between genetically different mice infected with *P. chabaudi*. (A-E) Change in health parameters over time in eight parental mouse strains infected with *P. chabaudi* (N= 5 A/J, N=5 C57BL/6J, N=5 129S1/SvlmJ, N=5 NOD/ShiLtJ, N=5 NZO/HILt J, N=4 CAST/EiJ, N=7 PWK/PhJ, N=5 WSB/EiJ). (F-J) Change in health parameters over time in DO mice (N= 489) and average change in parental mice without standard error. The health parameters monitored are as follows: (A, F) percent change in weight, (B, G) change in temperature, and (C, H) RBC count. All of these parameters decrease during the infection. On the contrary, (D, I) parasitemia (percent of infected RBC amont all RBC), and (E, J) parasite density (parasite load per mL of blood) both increase during malaria infection. All variables fluctuate at different rates and intensities during malaria infection.

We measured correlations between extreme symptom levels to determine how each disease sign correlated with each other. For example, we measured peak parasitemia, minimum RBC count, minimum temperature and minimum weight. When we plot RBCs, temperature or weight against maximum parasitemia we produce disease tolerance curves, which report how sick an individual will get in response to their parasite load. We found that the DO mouse disease outcomes largely recapitulate the spectrum of parental strain disease responses; DO mice stay within this phenotype envelope creating a continuous range of phenotypes (Figure 2). We found that RBC concentration and temperature varied with pathogen load but we did not see a correlation with weight changes and parasite load. The scatter plots are quite continuous and there aren’t population islands of mice that behave grossly different from the other mice.

**Figure 2.**
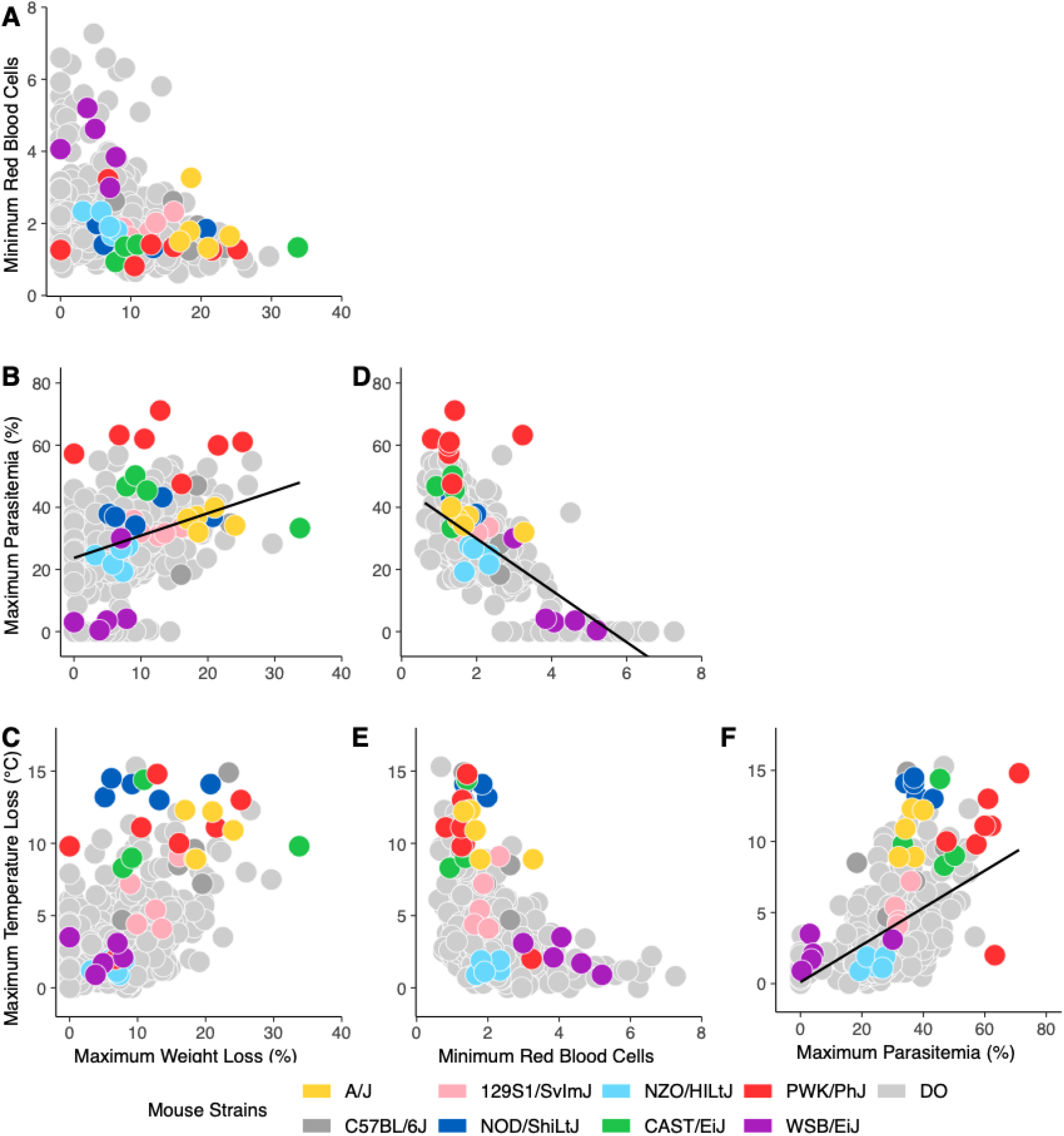
Tolerance curves mapping infection dynamics in parental mice strains and DO mice reveals a continuous range of phenotypes Multidimensional malaria infection health manifold showing DO mice and parental strain mice. The parameters monitored are as follows: (A, B, C) Maximum change in weight, (A, D, E) minimum red blood cell count, (B, D, F) maximum parasitemia, and (C, E, F) maximum change in temperature of infected DO mice (N= 489). Each health parameter is plotted against the others to obtain tolerance curves that relate extremes of health regardless of timing. R-squared for B, D, F are 0.1295, 0.5078 and 0.2984 respectively.

To check that our approach was working, we mapped recessive parental strain phenotypes for white coat color, black coat color, and a white forehead blaze that are segregating in the DO population and have known genetic causes (Figure S1) (Svenson *et al.* 2012). As anticipated, we mapped the white coat color phenotype (which presented in 90 of our 489 experimental mice) to the tyrosinase (*Tyr*) gene at 87.5 Mb (87.36-87.72 Mb, p<0.01) on chromosome 7 (Figure S1a). Our parental contribution analysis confirmed that A/J and NOD/ShiLtJ, both albino strains, were the contributors of this trait to the population. The black coat trait is caused by a recessive allele in *nonagouti* (*a*) on chromosome 2. We found 62 of our 489 experimental DO mice had black coats. We mapped this phenotype to a locus on chromosome 2 at 154.7 Mb (154.64-155.22 Mb, p<0.01), the site of the *a* gene. The parental contribution identified A/J and C57BL/6 as the two parental strains with recessive *a* alleles. Note that there is an interval from 76.9 Mb to 86.2 Mb on chromosome 2 where our analysis cannot map the WSB/EiJ parental contribution properly. This occurs because mice carrying WSB/EiJ specific traits in this interval were culled from the population by JAX in early DO generations (Figure S1b). A homozygous loss of function mutation in *Kitl* produces a white spot of hair on the forehead (blaze or star). We examined all of the non-white mice with the blaze (as it is not possible to determine if white-coated mice have a white forehead blaze) and mapped this trait to the previously identified locus on chromosome 10 at 89.6 Mb (89.57-89.62 Mb, p<0.01). This trait is introduced into the DO population by the WSB/EiJ parental strain (Figure S1c).

Previous studies identified *Char* loci that contribute to *P. chabaudi* disease severity in inbred strains using survival, parasitemia, parasite growth rate, parasite clearance, erythropoiesis, and more. We reasoned that these traits may also be found in the DO collection and therefore searched for the *Char* loci directly in our DO data, using the survival phenotype (Figure S2). *Char12*, found on chromosome 8, had the highest LOD score of the *Char* loci, though it did not reach statistical significance.

We asked whether readily quantifiable disease phenotypes correlated with SNPs in this population (Table S2). These phenotypes included simple continuous traits such as maximum parasite load or minimum RBC counts as well as more complex measurements such as rates of change or the timing of phenotypes. None of these continuous traits gave us a statistically significant result. We were able to map two statistically significant binary traits: no parasitemia and death (Table 1).

**Table 1.** Phenotypes and QTL intervals. ^1^Logarithm of the odds

| **PHENOTYPE** | **QTL REGION** | **LOD<sup>1</sup>** | **P-VALUE** | **FOUNDER EFFECT** | **FIGURE REFERENCE** |
| --- | --- | --- | --- | --- | --- |
| No Parasitemia | Chr 2:<br>79.30991-79.36893 Mb | 11.2<br>7571 | <0.01 | Undetermined | Figure 3A-C |
| Survival | Chr 10:<br>41.39297-43.77759 Mb | 8.72<br>6616 | <0.01 | 129S1/SvImJ increase | Figure 5A-C |

We found that 23 of the 489 infected DO mice had no detectable levels of parasites at any point during the infection (Figure 3A). We mapped a locus controlling this phenotype to a 0.59 Mb locus on chromosome 2 (p<0.01) (Figure 3B). The parental contribution in Figure 3B is obscured by a WSB/EiJ (purple) meiotic drive at this locus. The interval contains just two genes t, *Itga4,* an integrin subunit, and *Cerkl,* a ceramide kinase-like protein (Figure 3C). Note that this mapping reveals the same problem with gene mapping on chromosome 2 as we saw when mapping the blaze phenotype.

**Figure 3.**
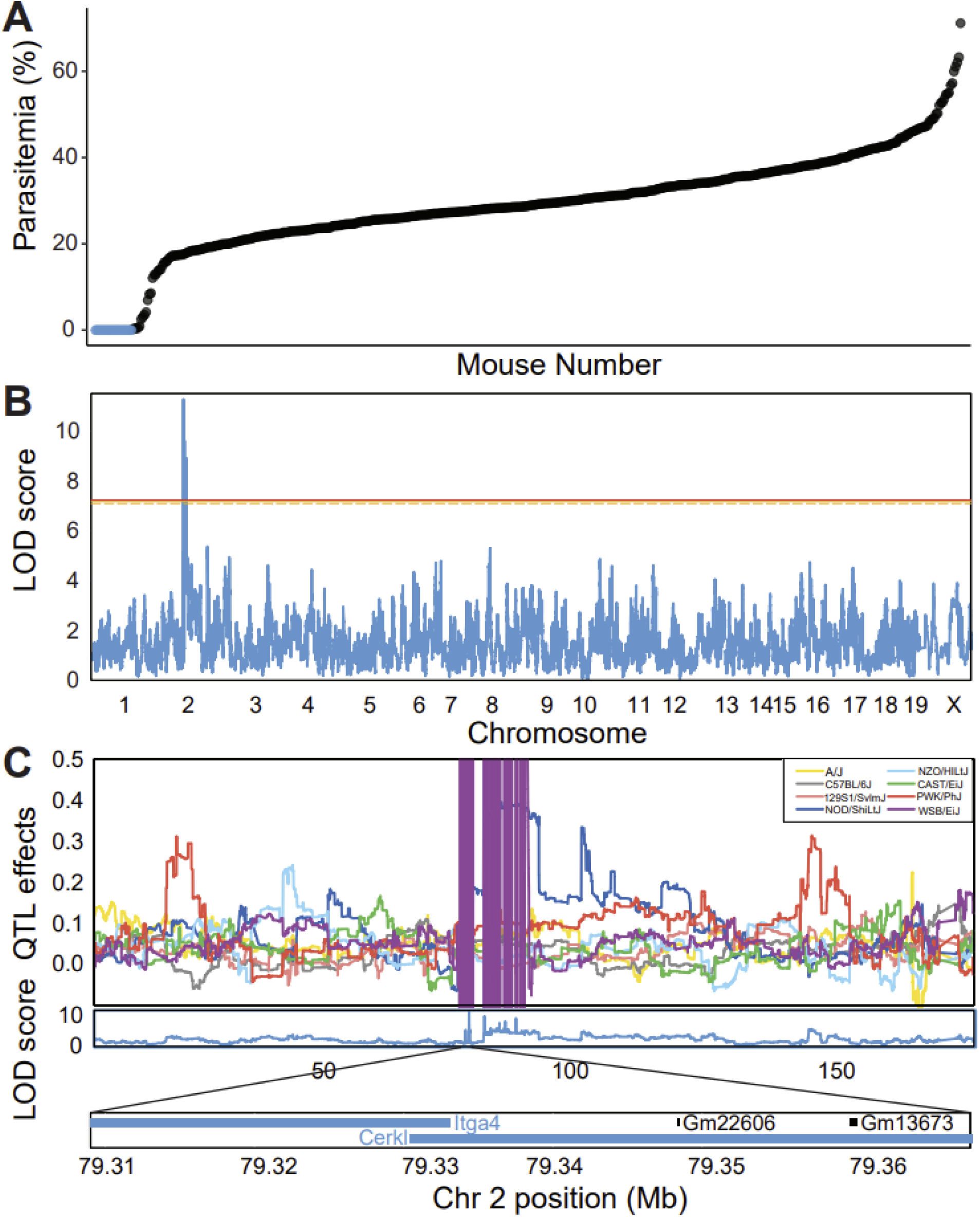
Undetectable Parasitemia Trait Maps to Chromosome 2 (A) DO mice infected with *Plasmodium chabaudi* AJ scattered based on peak percent parasitemia (N=489). 23 of the 489 experimental mice had no detectable parasitemia at any point at the infection (shown in blue). (B) Binary quantitative trait loci mapping for undetectable parasitemia. (undetectable parasitemia = 1, Others = 0) has a peak on chromosome 2. Horizontal lines show the significance threshold from permutation testing (p-value = 0.05 (orange line), 0.1 (orange dashed line)). (C) Parental contributions for undetectable parasitemia trait loci on chromosome 2 were obfuscated by the presence of *R2d2* meiotic drive from the WSB/EiJ parental strain that was removed from the DO population.

**Figure 4.**
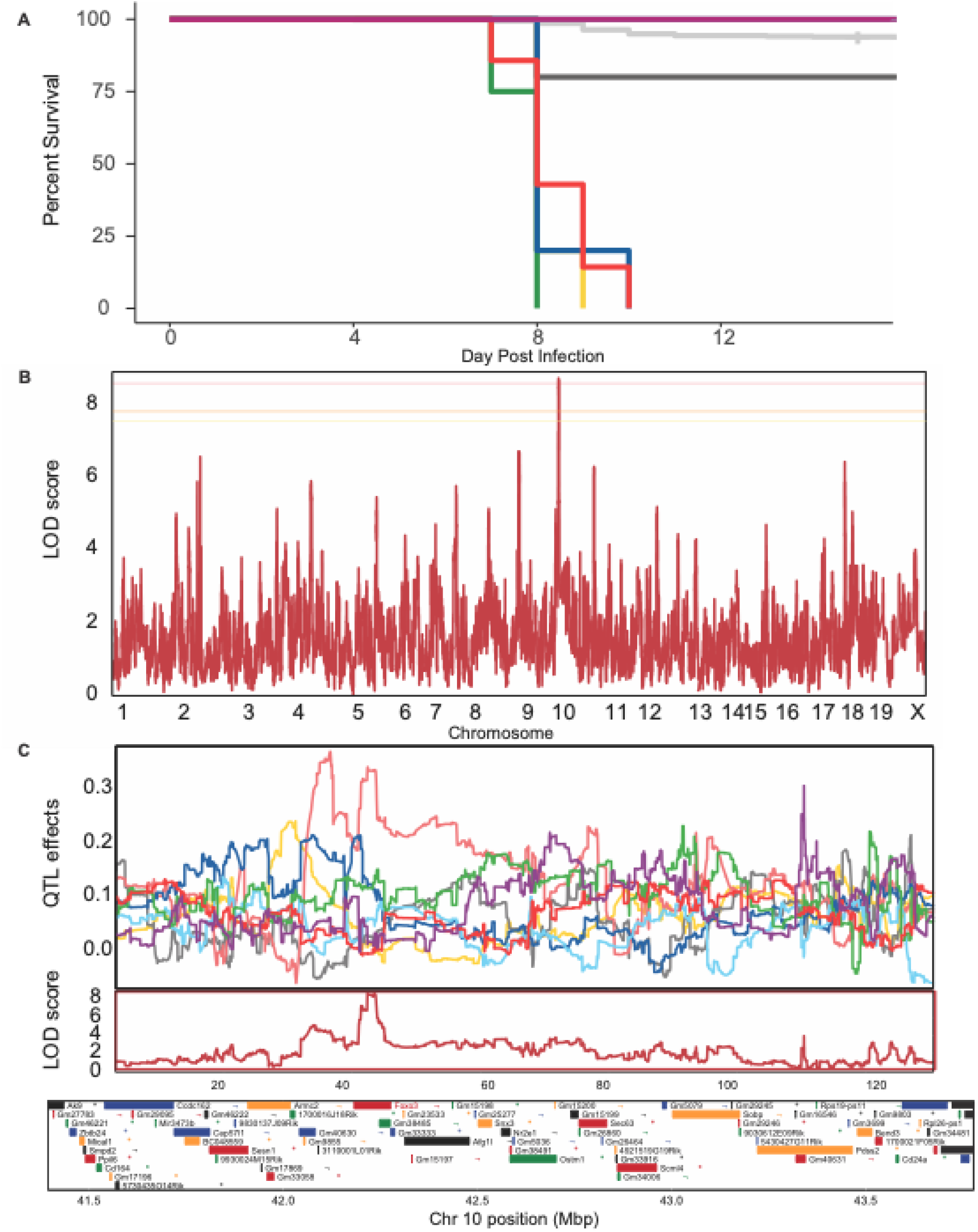
Survival Trait Maps to Chromosome 10 (A) Kaplan-Meier plot of collaborative cross parental strains (N= 5 A/J, N=5 C57BL/6J, N=5 129S1/SvlmJ, N=5 NOD/ShiLtJ, N=5 NZO/HILt J, N=4 CAST/EiJ, N=7 PWK/PhJ, N=5 WSB/EiJ) and DO mice (N=489) infected with *Plasmodium chabaudi* AJ. CAST/EiJ, A/J, NOD/ShiLtJ, and PWK/PhJ succumb to infection. C57BL/6J and the DO mice were partially resistant to infection. WSB/EiJ, NZO/HILtJ and 129S1/SvlmJ were resistant to infection. (B) Binary quantitative trait loci mapping for *P. chabaudi* infection survival (Died =1, Lived = 0) has a peak on chromosome 10. Horizontal lines show the significance threshold from permutation testing (p-value = 0.01 (red line), 0.05 (orange line), 0.1 (orange dashed line)). (C) Parental contribution to survival trait locus on chromosome 10 with candidate gene *Foxo3* discussed in text highlighted in indigo. (D) DO mice (N=489) infected with *Plasmodium chabaudi* AJ clustered in mild (83 mice), moderate (379 mice), and severe (27 mice) (severe cluster shown in red). (E) Binary quantitative trait loci mapping for severe parasitemia. (Severe parasitemia = 1, Others = 0) has a peak on chromosome 10. Horizontal lines show the significance threshold from permutation testing (p-value = 0.05 (orange line), 0.1 (orange dashed line)). (F) Parental contributions for severe parasitemia trait loci on chromosome 10 candidate gene *Foxo3* discussed in text highlighted in indigo.

The second locus we found by mapping the mice that didn’t survive the infection. We found the parental strains exhibited a broad phenotypic range from nonlethal (WSB/EiJ, 129S1/SvlmJ, NZO/HILtJ), to partial lethality (C57BL/6), to inevitable lethality (NOD/ShiLtJ, A/J, PWK/PhJ, CAST/EiJ). The DO experimental mice show higher survival to *P. chabaudi* challenge compared to the susceptible parental strains; 29 of the 489 mice died (Figure 5A). Mouse mortality was limited to days 7-10 days post-infection; mice that survived that time interval recovered. We identified a significant 2.38Mb locus on chromosome 10 (p<0.01) (Figure 5B). 129S1/SvlmJ provided a high parental contribution in this region (Figure 5C). This interval contains multiple gene coding regions with SNP variants among the eight parental strains, including *Foxo3,* an immune modulator in monocytes (Figure 5C).

## Discussion

We explored the range of disease phenotypes found in a genetically diverse population of mice in response to *Plasmodium chabaudi chabaudi* AJ infection. We studied DO mice and their parental strains to reveal the range of phenotypic variation by taking longitudinal measurements of weight, temperature, anemia, parasite loads and survival. Figure 1 shows the breadth of phenotypes for each of these parameters. In general, the DO mice were more resilient to infection compared to their parental strains as seen by the greater proportion of the DOs that survived the infection. Perhaps the inbred parental strains are more likely to express extreme phenotypes because they are homozygous for many recessive traits. The continuous distribution of malaria phenotypes found in DO mice supports the claim that the host response is polygenic, which is also suggested to be the case in human populations (Damena and Chimusa 2020).

We found that it was possible to make significant disease tolerance curves that correlated temperature loss, RBC loss and weight loss with maximum parasite loads. The R^2 values ranged from 0.5078, 0.2984 and 0.1295, respectively, for temperature, RBCs and weight versus parasite load. None of these quantitative traits was mappable to a small number of loci.

We tested a large variety of disease phenotypes that we could extract from our dataset (Table S2) and two binary traits gave us statistically significant hits that can be starting points for further exploration.

We could not detect parasites in 23 of the infected DO mice. This phenotype correlated with a 0.6Mb interval on chromosome 2 containing two genes (*Itga4*, *Cerkl*). *Itga4* encodesthe alpha 4 subunit of an integrin family of proteins that play a role in cell motility and migration. This integrin is a therapeutic target for the treatment of multiple sclerosis and inflammatory bowel disease in humans *(Gordon *et al.* 2002; O’Doherty et al. 2007; Soler et al. 2009)*. *Cerkl* codes for a ceramide kinase-like protein (Cerkl) and was found to maintain mitochondrial thioredoxin in a reduced redox state. Through this interaction, Cerkl plays a role in protecting cells from apoptosis under oxidative stress (Li *et al.* 2014). In humans, mutation of this gene was shown to cause autosomal recessive retinitis pigmentosa (Tuson *et al.* 2004). These genes have not been implicated in microbial pathogenesis previously and their role is not immediately obvious.

This region on chromosome two presents mapping problems in the DO mice collection. A meiotic drive element from the WSB/EiJ strain was found to be replacing all other parental contributions at the *R2d2* locus from 76.9 Mb to 86.2 Mb on chromosome 2 (Didion *et al.* 2015) in the DO population. To protect the diversity of the DO population in this interval, researchers at JAX performed an artificial selection against mice with the WSB/EiJ allele in the *R2d2* region from the DO population (Chesler *et al.* 2016). This is problematic for us because this locus is linked to our zero parasitemia trait, which we reason comes from the WSB/EiJ mice and mapped to this region. The QTL analysis software we used fails when mapping chromosome contributions when one of the eight parental strains is missing from the cross, which results in the extreme positive and negative LOD scores for WSB/EiJ in this region.

We mapped a 2.38 Mb interval on chromosome 10 correlating with death by malaria. This interval contains 20 coding genes with SNP variants among the eight parental strains. One of the genes contained in the interval, *Foxo3,* an immune modulator in monocytes, has known non-coding polymorphisms that are significantly associated with malaria severity in humans (Lee *et al.* 2013; Nguetse *et al.* 2015). *Foxo3* drives an inflammatory response in monocytes which reduces pro-inflammatory cytokines TNF-α and increases anti-inflammatory cytokines like IL-10 in a TGFβ1-dependent manner. In addition to this known coding gene, the identified locus of interest includes many uncharacterized proteins that could be relevant to the infection and worth further investigation.

Previous QTL analyses describing resistance against *P. chabaudi* infections identified ten *Char* genes using inbred cross studies (Huang *et al.* 2018). We identified *Char12* on chromosome 8, although as an insignificant peak (Figure S2). Other previously identified *Char* genes were not discernable against the baseline noise in the QTL analyses. It is possible that there was no variation of *Char* alleles in this population, which would make them impossible to find in our experiment. We also used a different strain of Plasmodium (AJ versus AS) than had been used in previous studies, and it is possible that the Char phenotype is parasite strain specific.

Why is it that we were successful in mapping binary traits for malaria disease but not any of the quantitative phenotypes we observed? The good news about this study is that these quantitative phenotypes are variable which means that there is hope that we can learn to manipulate them. It is frustrating, however, that none of these were mappable. One possible explanation is that the quantitative infection traits are affected by a large number of loci, and we don’t have the power to map these traits using ~500 mice. Put another way, perhaps we are following phenotypes that matter to scientists, but not to the mice: For example, we picked easy phenotypes to measure, but something like temperature or weight loss could conceivably involve many inputs. This leaves open the question about how we choose the phenotypes we decide to measure.

## Author Contributions

ASG, VC, and DSS conceived and designed the experiments. ASG and VC performed the experiments. ASG, VC, ASK, and DSS analyzed the data. ASG, VC, ASK, and DSS wrote the paper with contributions from NMD, KC, ML.

## Acknowledgments

The authors would like to thank the Department of Microbiology and Immunology at Stanford for being supportive of our research and professional development. We extend an additional thank you to Denise M. Monack, Liliana M. Massis, Katherine Cumnock, and Michelle Lissner. In addition, Karl Broman and Daniel Gatti were gracious advisors in putting together the analysis. Finally, the authors would like to thank the Affymetrix Life Sciences Bioinformatics Team for all the help and support in the genotyping array. This study was supported by a contract from the Defense Advanced Research Projects Agency (W911NF-16-0052) (D.S.S.).

## Supplemental Information

**Table S1.** Mice Used in Experiment.

| MOUSE STRAIN | STOCK NUMBER |
| --- | --- |
| J:DO | 009376 |
| A/J | 000646 |
| C57BL/6J | 000664 |
| 129S1/SvImJ | 002448 |
| NOD/ShiLtJ | 001976 |
| NZO/HILtJ | 002105 |
| PWK/PhJ | 003715 |
| CAST/EiJ | 000928 |
| WSB/EiJ | 001145 |

**Table S2.**
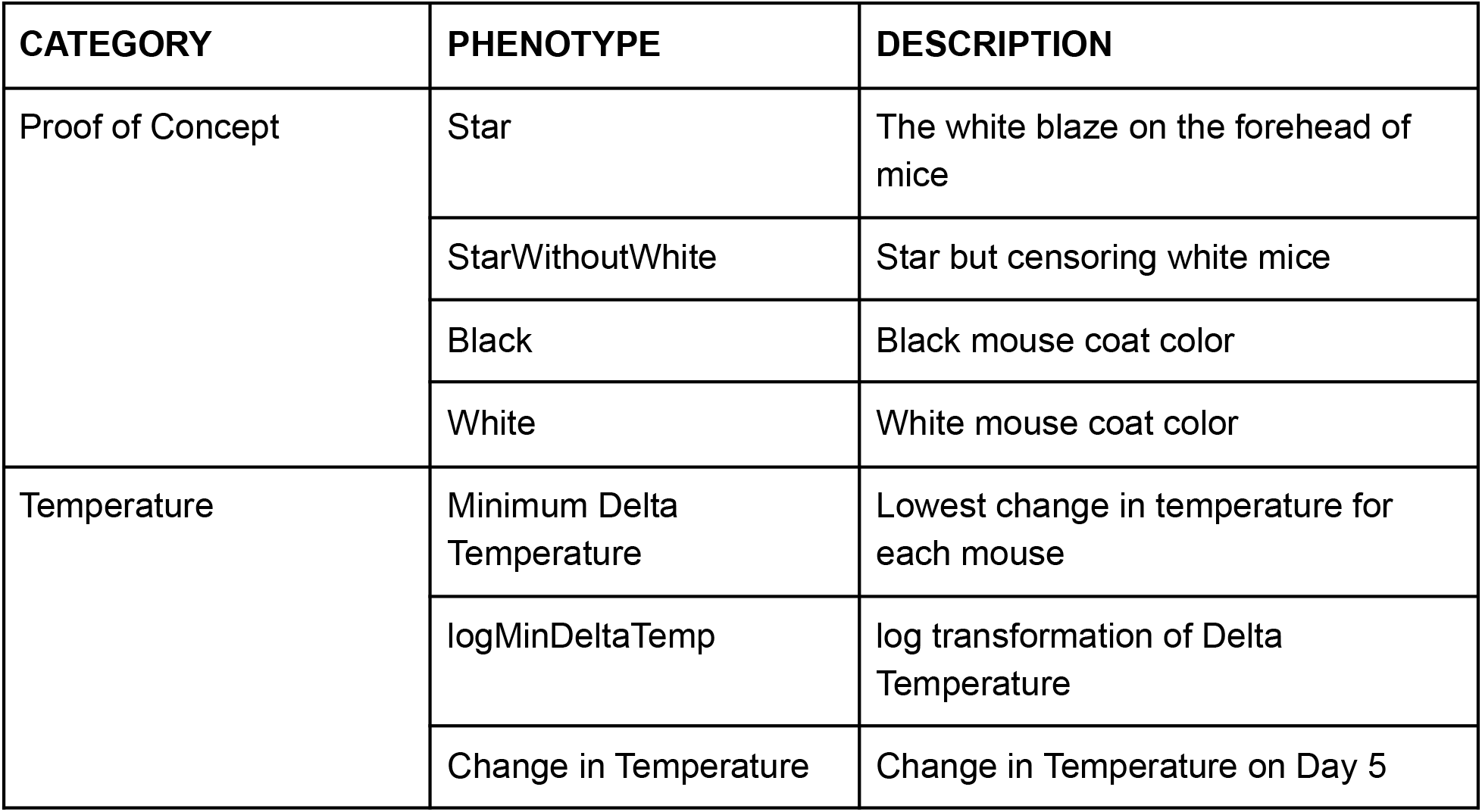
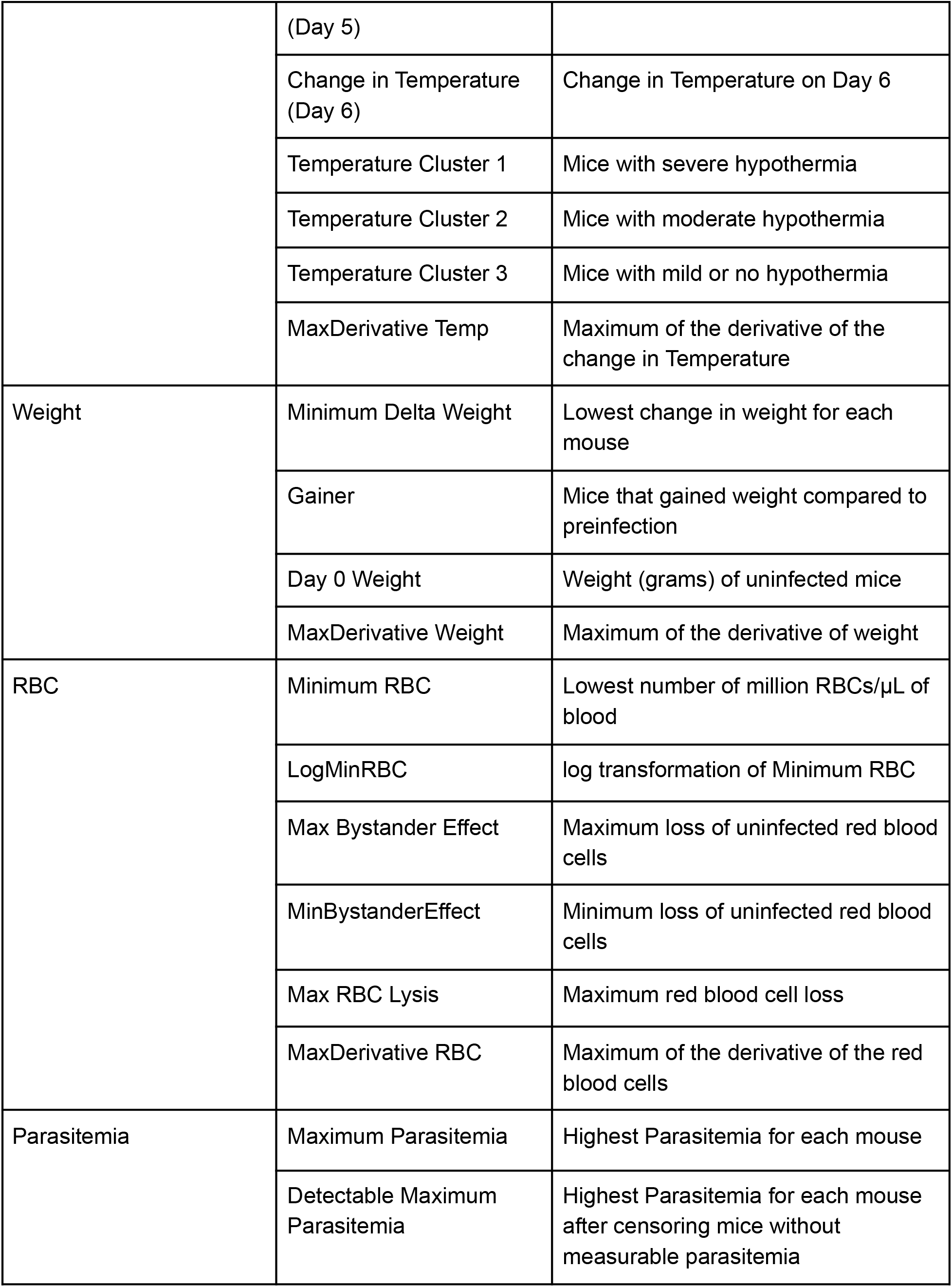
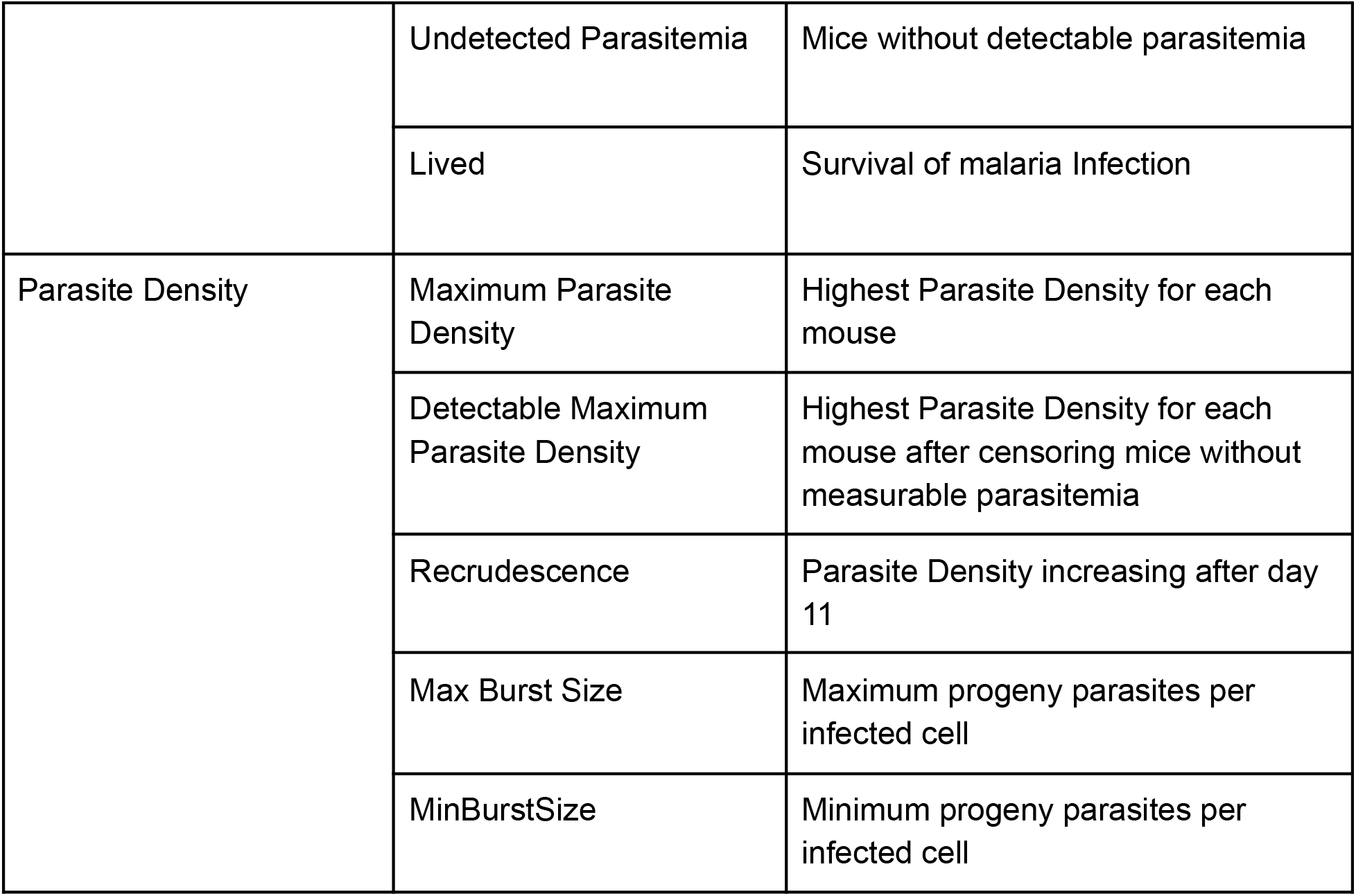
List of phenotypes

Table S3. Full sampling sheet

**Table S4.**
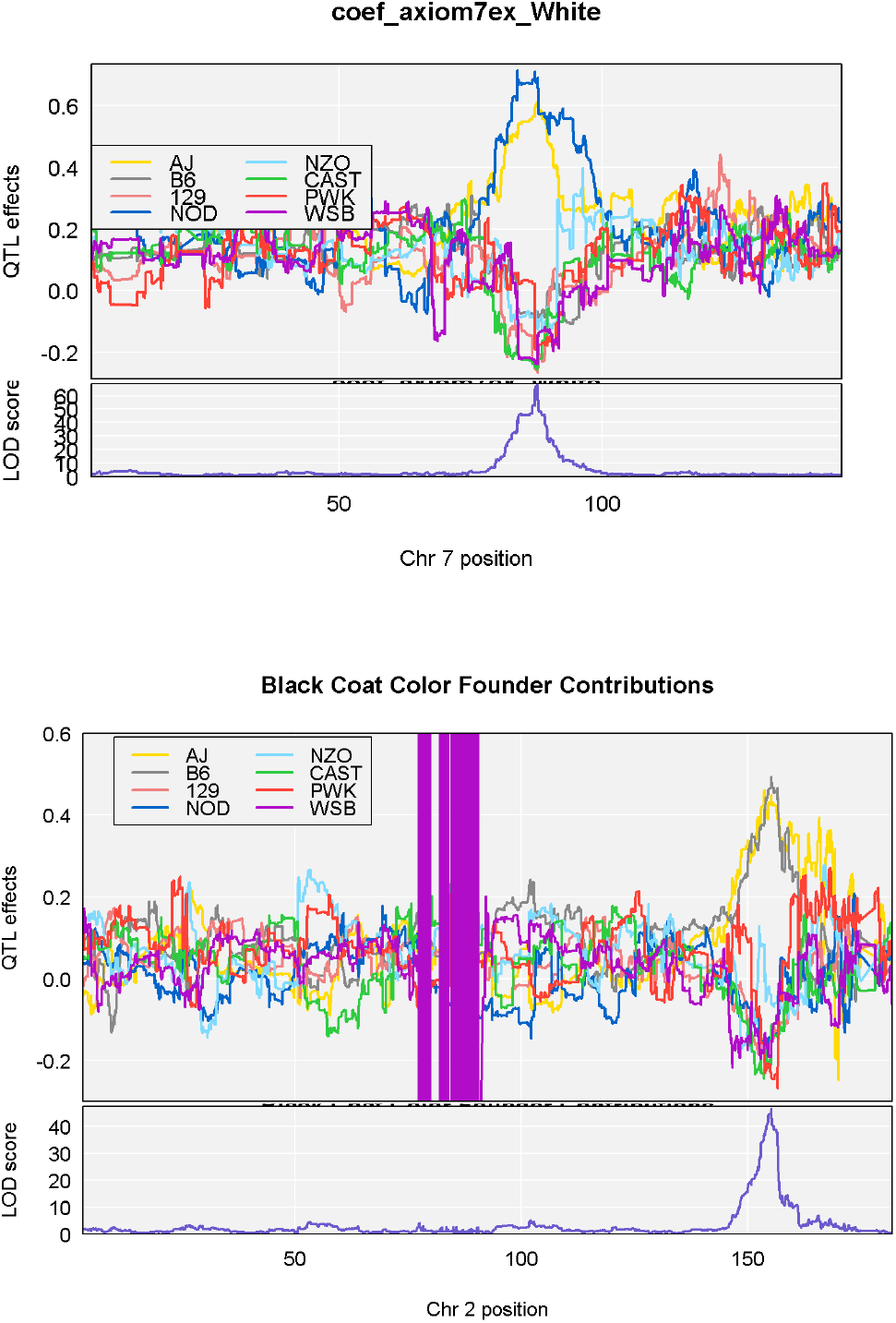
Full genotyping data matrix

**Figure S1.**
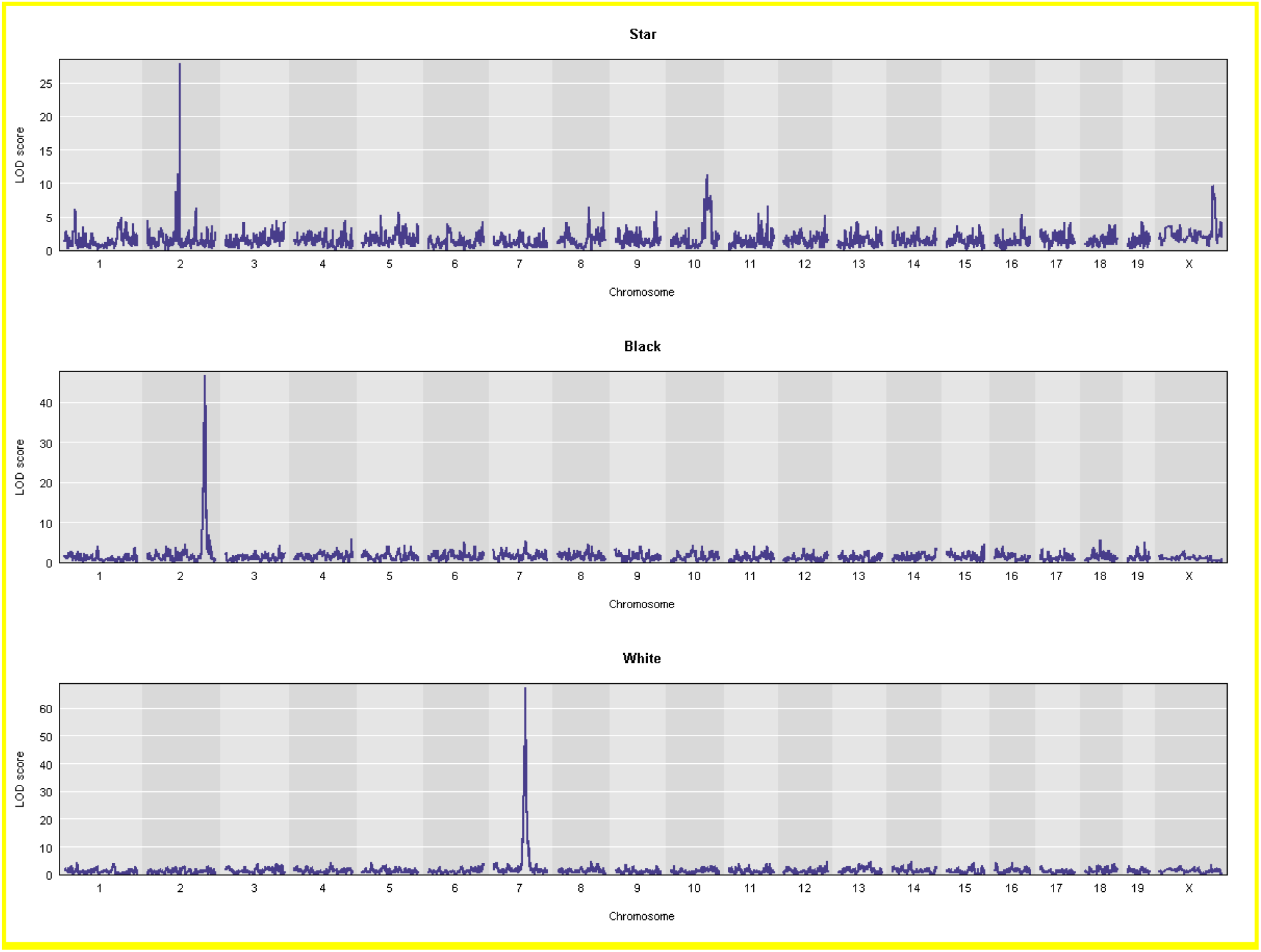
Genome-wide scan for coat color traits as proof of concept for QTL analysis A and C) The LOD at each SNP from the eight state linear model. Colored lines show permutation derived significance thresholds at p = 0.05 (red), p=0.10 (orange), and p=0.2 (yellow). B and D) Founder strain effect plots for each phenotype on the chromosome with the maximum QTL.

**Figure S2.**
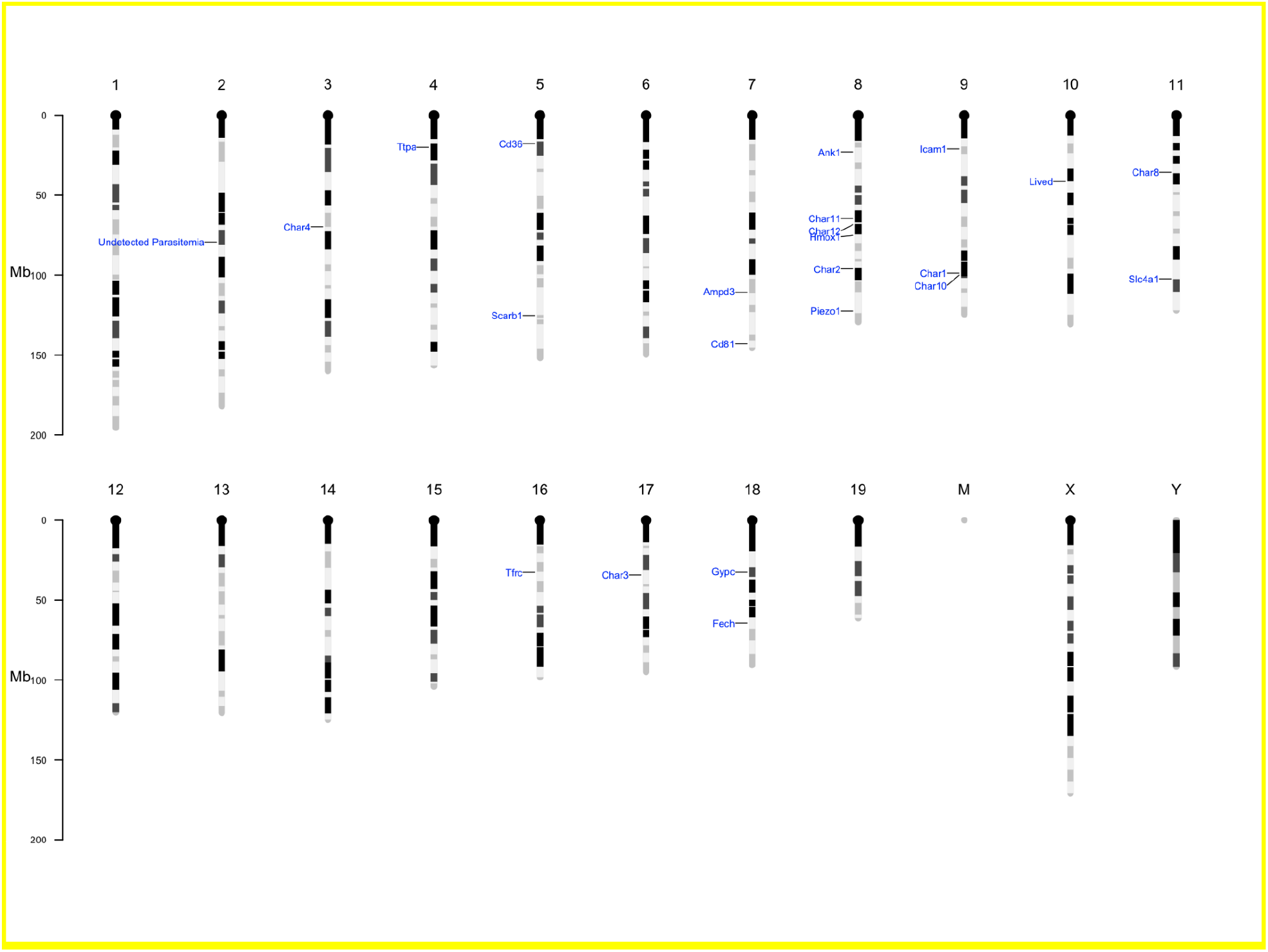
All Chars using survival phenotype

**Figure S3.**
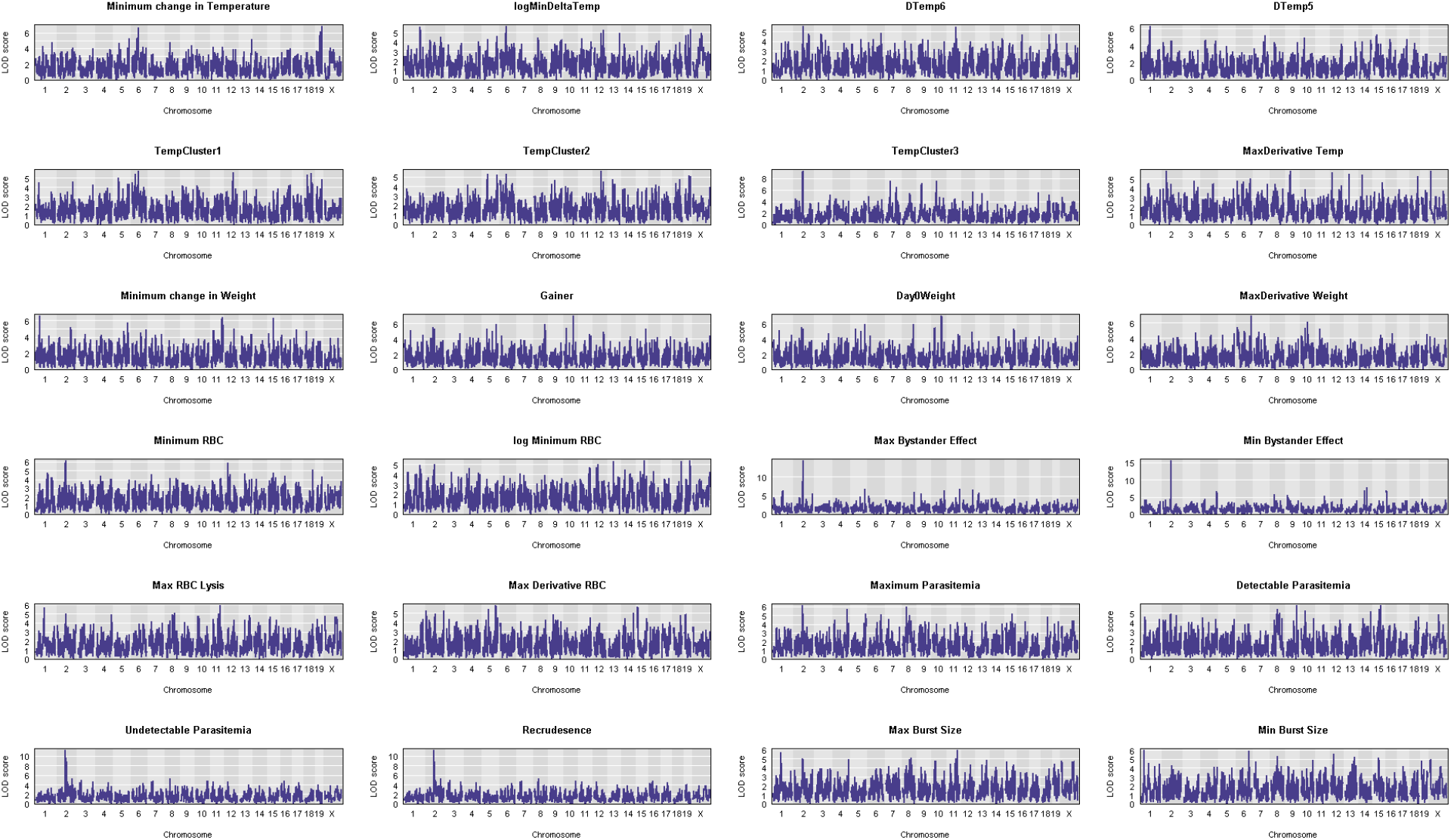
QTLs that did not meet significance threshold

## Notes

### Competing Interest Statement

The authors have declared no competing interest.

